# Pupil responses to hidden photoreceptor– specific modulations in movies

**DOI:** 10.1101/440040

**Authors:** Manuel Spitschan, Marina Gardasevic, Franck P. Martial, Robert J. Lucas, Annette E. Allen

## Abstract

Under typical daytime light levels, the human pupillary light response (PLR) is driven by the activity of the L, M, and S cones, and melanopsin expressed in the so-called intrinsically photosensitive retinal ganglion cells (ipRGCs). However, the importance of each of these photoreceptive mechanisms in defining pupil size under real-world viewing conditions remains to be established. To address this question, we embedded photoreceptor-specific modulations in a movie displayed using a novel projector-based five-primary spatial stimulation system, which allowed for the precise control of photoreceptor activations in time and space. We measured the pupillary light response in eleven observers, who viewed short cartoon movies which contained hidden low-frequency (0.25 Hz) silent-substitution modulations of the L, M and S cones (no stimulation of melanopsin), melanopsin (no stimulation of L, M and S cones), both L, M, and S cones and melanopsin or no modulation at all. We find that all photoreceptors active at photopic light levels regulate pupil size under this condition. Our data imply that embedding modulations in photoreceptor contrast could provide a method to manipulate key adaptive aspects of the human visual system in everyday, real-world activities such as watching a movie.

## Introduction

Photoreception in the human retina proceeds from rods, cones and the photopigment melanopsin, which is expressed in a subset of so-called <u>intrinsically photosensitive retinal</u> <u>ganglion cells</u> (ipRGCs) [1–5]. All retinal photoreceptors contribute to visual function, though their exact contributions in naturalistic behaviour depend on the spatial and temporal characteristics of the retinal stimulus [6]. At photopic light levels, pupil size is controlled by an excitatory input of L+M (luminance), an inhibitory input of S cones (and possibly M cones, see [7, 8]), as well as a positive input from melanopsin (Fig. 1A), along with small input from rods. Since the pupil, the aperture of the eye, changes its size depending on the activity of the photoreceptors in the retina, pupillometry represents a non-invasive and convenient method of assessing photoreceptor function.

**Figure 1.**
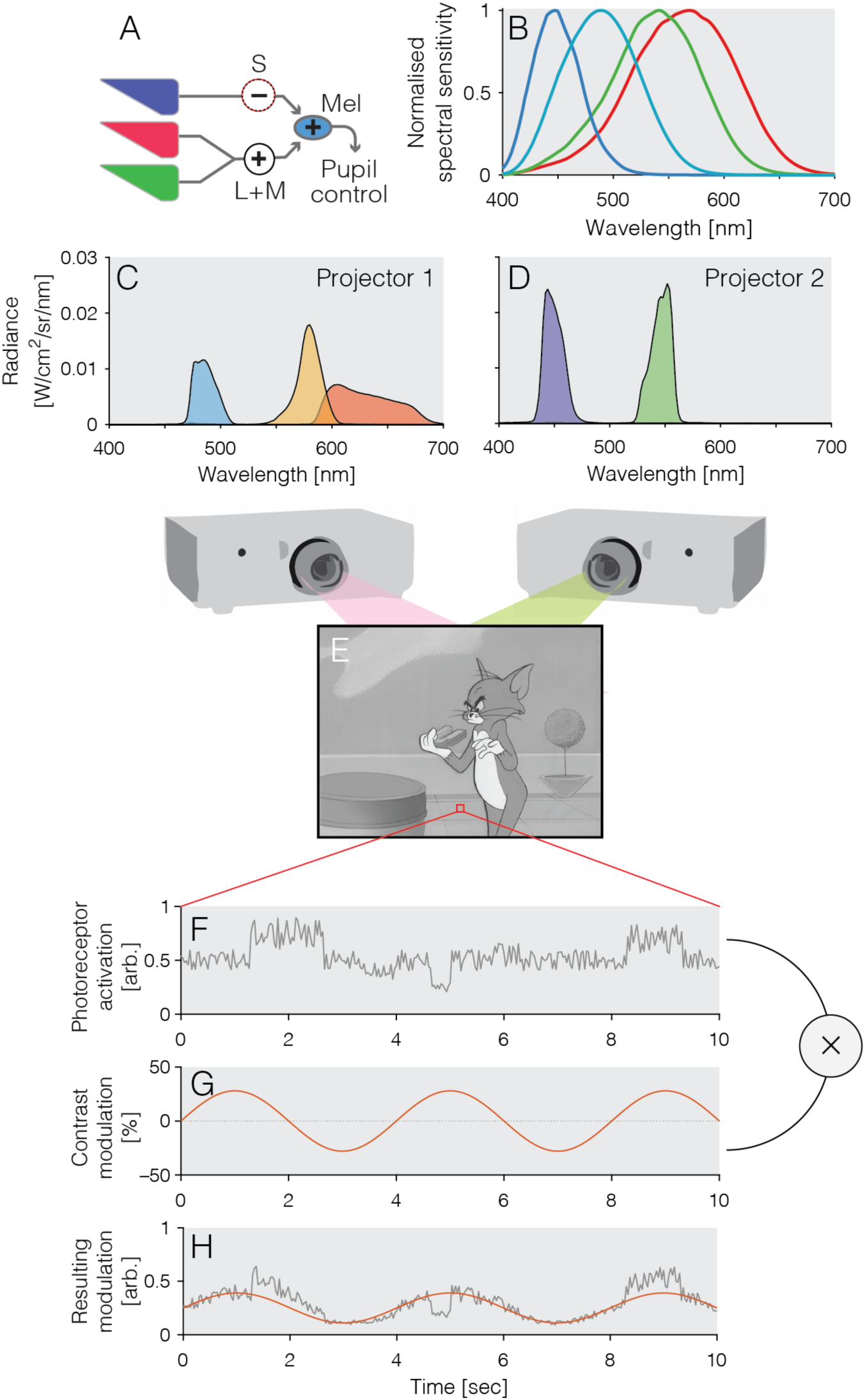
Fundamentals and methods. *A*) Spectral sensitivities of the human photoreceptors; *B*) Schematic circuit diagram of the human pupillary light response, with L + M cones, and melanopsin, providing an excitatory drive to pupil constriction, and S cones yielding an inhibitory input; *C and D*) Spectral radiance of the five primaries for the two projectors; *E*) Sample frame from the Tom & Jerry cartoon; *F*) Sample time course over 10 sec of one pixel of the cartoon, displaying local variations in luminance; *G*) Time course of a pure sinusoidal stimulus (0.25 Hz) stimulating a given photoreceptor class; *H*) The sinusoidal stimulus get superimposed with the variations in pixel luminance so as to yield an embedded sinusoidal modulation while retaining image contrast.

The contributions of different photoreceptors to pupillary control have been characterised in analytical and parametric studies under well-controlled but rather reduced stimulus conditions typically employing spatially homogenous fields of different visual extent [7–16]. However, under real-world conditions, the importance of each photoreceptor class in defining pupil size remains to be established. In this study, we examined the possibility to address this question by embedding photoreceptor-specific modulations into movies seen under real-world viewing conditions.

In this work, we used a modified projector system, the design of which was described previously [17], to examine the pupil responses to photoreceptor-selective sinusoidal modulations embedded in short greyscale Tom & Jerry cartoon clips. Under photopic conditions, we examined responses to stimuli targeted at the *LMS* cones (with no modulation of melanopsin), *melanopsin* (with no modulation of LMS cones), and a modulation of all photoreceptors to the same extent, which we term *Light Flux.* We find evidence that all photoreceptors regulate pupil size to some degree even during high spatiotemporal frequency modulations.

## Methods and materials

### Visual stimuli

Stimuli were presented in free-viewing, and participants were free to move their eyes throughout the study. Participants were seated 128 cm from a canvas onto which two spatially aligned projectors projected a movie which contained an embedded 0.25 Hz modulation of photoreceptor contrast enabled by the method silent substitution (see below).. The stimulus was a rectangle of dimensions 81 × 109 cm, yielding a total visual angle of approximately 35 × 46 deg2. The targeted photoreceptors were the LMS cones at equal contrast (*LMS*), melanopsin (*Mel*), or all photoreceptors in unison (*Light Flux*). A fourth condition (*Reference*) contained no modulations. In our low-frequency modulations, we did not control for the effect of cones in the shadow of the retinal blood vessels, so-called penumbral cones, which are known to be stimulated in high-frequency (8-16 Hz) melanopsin-directed modulations [18].

### Principle of silent substitution

The spectral tuning of the individual photoreceptors is relatively broadband, and therefore any given light will activate all photoreceptors (Fig. 1B). Measurement of downstream responses, such as pupil size, using narrowband, or monochromatic light, will therefore only yield non-specific responses. For example, while in the living human eye, melanopsin will respond maximally to a monochromatic ∼488 nm light (the shift from 480 nm of peak pigment absorbance to longer wavelengths due to pre-receptoral filtering by the lens and ocular media), such a light will also activate S cones at ∼41%, M cones at ∼43%, and L cones at ∼26% of their maximal response, respectively. This non-specificity can be overcome by the method of silent substitution [19, 20]. In this method, pairs of spectra are presented to an observer such that the exchange between the spectra only stimulates one class of photoreceptors (the *stimulated* photoreceptors), while not changing the activation of the set of *silenced* photoreceptors. Silent substitution can be achieved using any light source which has at least as many independent lights as spectral photoreceptors under examination, such as sets of LEDs combined optically [7, 9–11, 13–16, 21–23], or combination of broadband light filtered through a diffraction grating and imaged on digital micromirror device [12, 18, 24, 25] (see [26] for technical details), or using modified projectors [17, 27] as in this study.

### Projector system

The design of the projector system was first described in [17], but this concrete realization was slightly different. Two 8-bit Hitachi CP-X5022WN 3LCD projectors were stacked on top of each other and the boundaries of their respective projected outputs were aligned by the experimenters using lens adjustment in vertical and horizontal axes, which was further refined with digital cornerstone correction. Alignment was confirmed by visual inspection of corners and edges and in subsequent experiments (not reported here) using high-spatial frequency stimuli. The projectors were modified using optical filters. In Projector 1, the blue primary was modified using a 470 nm cut-off yellow longpass filter (PIXELTEQ, Largo, FL; part # LP470-r40×25×1), making a cyan primary; the green primary was modified using a 463-571 nm notch filter (MidOpt, Palatine, IL; part number #102340892), making yellow. In Projector 2, the blue primary was modified using a 463-571 nm magenta notch filter (MidOpt, Palatine, IL; part #102340892), making violet; and the green primary was modified using a 550 nm bandpass filter (PIXELTEQ, Largo, FL part # Bi550-r40×25×1), producing a narrower green primary. This yielded a dual projector system with five independent primaries (Fig. 1C, D). At maximum output, the primaries had CIE 1931 *xy* chromaticities: (0.65, 0.34), (0.50, 0.49), (0.08, 0.22), (0.28, 0.70), and (0.16, 0.02), and photopic luminances 54.36, 131.64, 20.84, 185.66, and 8.18 cd/m2, respectively. The projectors were calibrated and linearised using a CRS SpectroCAL spectrometer.

### Movie stimuli

The movie selected was the Tom & Jerry cartoon “Pent-House Mouse” (1963) [28] (Fig. 1E, length 6m11s). The film was first turned into a greyscale movie and then processed in MATLAB to embed photoreceptor-directed modulations into the luminance variations (Fig. 1F, G, H). The photoreceptor contrast of the modulation was 28% for all directions. The modulations were embedded as follows. The (assumed) linear intensity for pixel at position *x, y* at time *t* is given by *I*(*x, y, t*). The linear 0-1 value for primary *n* (where *n* ≤ 5) is then given by:

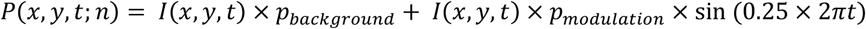
The resultant linear values per primary were then converted into 8-bit resolution.

### Photoreceptor spectral sensitivities

For the LMS cones, we assumed the physiologically-relevant CIE 2006 photoreceptor sensitivities [29]. Melanopsin spectral sensitivity was derived using the Govardovskii nomogram with lmax = 480 nm, assuming low photopigment density, and applying the same pre-receptoral filtering as for the cones. The receptor spectral sensitivities are available in the Silent Substitution Toolbox (http://github.com/spitschan/SilentSubstitutionToolbox).

### Age adjustment

To account for age-dependent differences in pre-receptoral filtering (specifically age-dependent lens-density), stimuli were adjusted in age group for each observer, with discrete categories 20-24 years of age, (22 years reference age) 25-29 years of age (27 years reference age), and 30-34 years of age (32 years of age). Because of the relatively large variability of lens density even across individuals of the same age [30, 31], we reasoned that adjusting according to age group in this way is an acceptable strategy. The age adjustment of pre-receptoral filtering was performed following the calculations set by the CIE [29]. Within those age groups, lens density does not abruptly change.

### Protocol and procedure

After giving informed consent, observers were tested for visual acuity and normal colour vision. Sitting down, they then viewed the four video conditions consecutively, with a two-minute break in between each video. The order of the conditions was randomized between participants. All testing took place during regular daytime working hours (9 to 5 pm).

### Pupillometry

The participants’ pupil responses were measured using a light-weight binocular eye tracking headset (Pupil Labs GmbH, Berlin, Germany), recording pupil responses at 30 Hz, and also a front-facing view of the participant’s view. Using software provided by the manufacturer, 2D ellipses were fit to the resulting videos off-line, and time-locked to the onset of the stimuli using open-source software provided by the manufacturer (https://github.com/pupil-labs/pupil).

### Participants

Eleven observers (mean age±SD 24.5±6.5 years, 7 female) were recruited from the wider University of Manchester community. Observers were entered into a raffle to win a hamper. Two participants were authors of this study (A.A. and M.G.). All subjects had 6/5 or better visual acuity (Snellen chart) and normal red-green colour vision as assessed by the Ishihara test [32].

### Data analysis

Raw pupil data were processed using custom MATLAB software as follows. Missing data reported from the 2D fitting algorithm were interpreted as eye blinks; to account for transient mis-estimations of the pupil size before and after eye lid closure, we removed additional samples around these events (padding window ± 5 samples at 30 Hz). These missing samples were then linearly interpolated. Data were them smoothed with a Savitzky-Golay filter (11th order, frame length 21 samples) using MATLAB’s built-in sgolayfilt() function. Raw pupil data (expressed in pixels of the 2D ellipse) were mean-normalised across the entire time series and expressed as percent change around that mean. We then extracted the per-cycle average pupil responses, collapsing across all cycles during the entire cartoon. This per-cycle average was then fit, in a least-squares sense, with a sum of sine and cosine at the stimulation frequency (0.25 Hz); the weights of the sine and cosine were then transformed into amplitude and phase of the evoked pupil response.

### Ethics

Ethical approval was obtained from the University of Manchester Ethics commission (approval number #2017-2958-43594359).

### Code and data availability

All data and code to perform the data analyses reported here can be found at https://github.com/spitschan/SpitschanEtAl2019_PupilMovies.

## Results

### Photoreceptor-selective modulations embedded in movies produce distinct pupil responses

The 0.25 Hz photoreceptor-specific modulation stimuli evoked distinct responses in all observers (Fig. 2A-D). The strongest response (in amplitude expressed as percent change in pupil diameter relative to the mean) was elicited by the *Light flux* modulations (mean±1SD = 5.0616±1.2554%, median = 5.3051%), followed by *L+M+S* (mean±1SD = 4.1340±0.9263%, median = 4.6839%) (Fig. 3A). In many observers we find a small melanopsin-evoked pupil response and in aggregate, there is a significant difference between the *Melanopsin* (mean±1SD = 2.1182±1.3096%, median = 1.5605%) and the *Reference* condition (mean±1SD = 1.0792±0.7652, median = 0.7462%) (Wilcoxon rank sum test, *Z* = 2.3639, *p* = 0.0181). We note that the melanopsin-evoked pupil response is somewhat inconsistent across observers, raising the possibility of individual differences in the observers’ melanopsin sensitivity (Fig. 2B; 3A, B).

**Figure 2.**
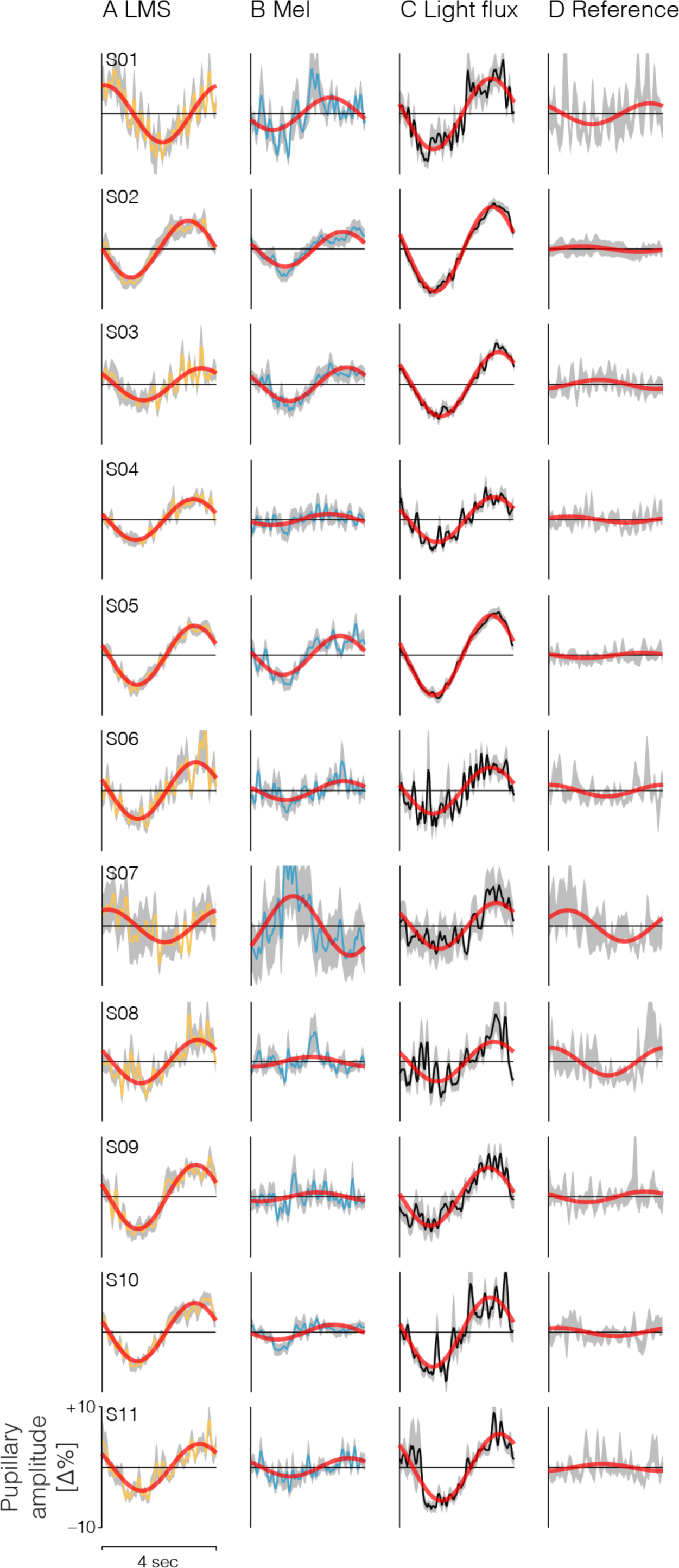
Individual subject pupil data. Data are collapsed (column-wise) across stimulus types. A) L + M+S, B) Melanopsin (*Mel*), C) Light flux (all photoreceptors are modulated), D) no modulation. Red lines indicate the least-squares fit of a sum and cosine at the modulation frequency to cycle-average response.

**Figure 3.**
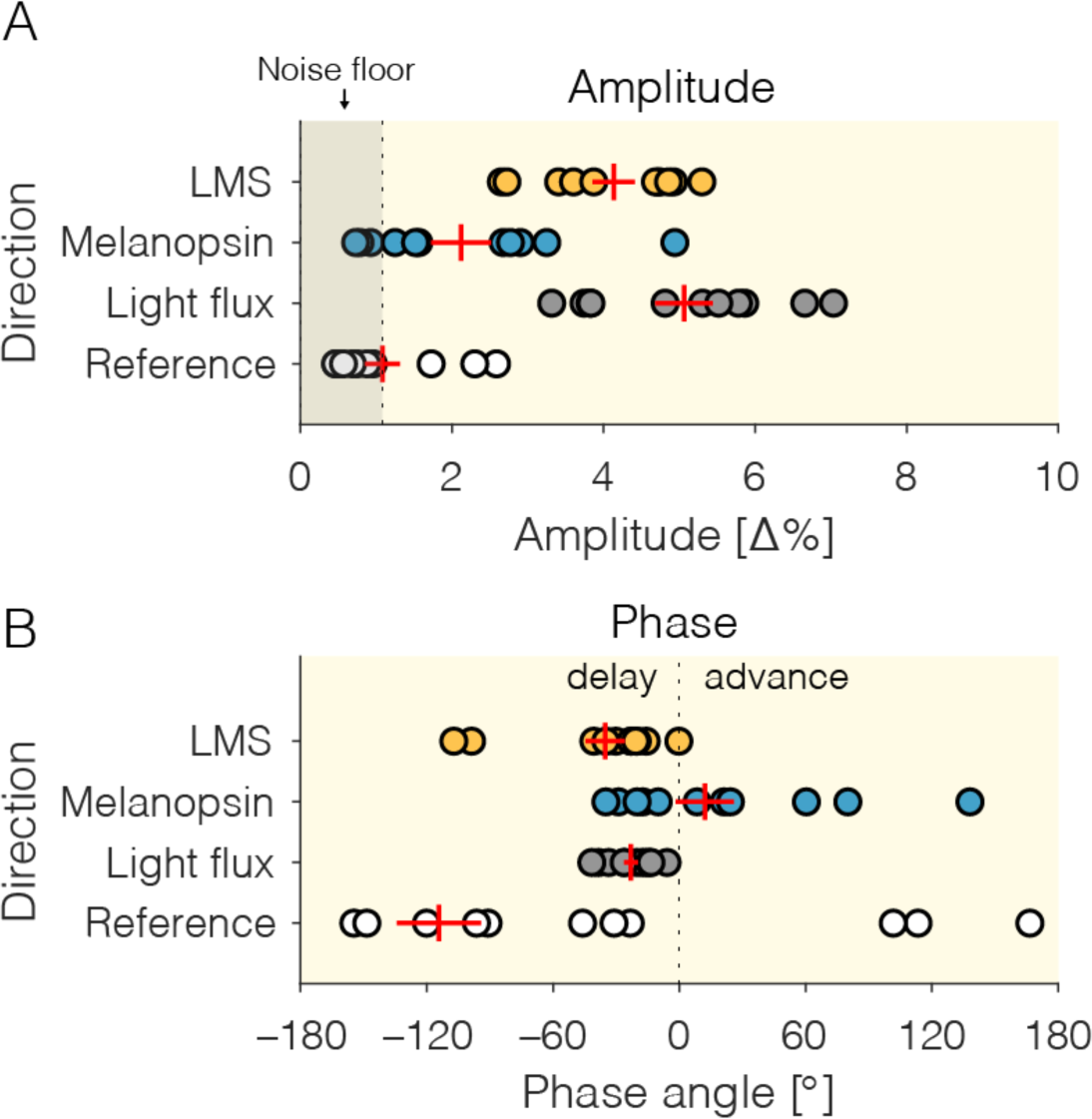
Extracted response properties of the pupil responses to photoreceptor-directed modulations embedded in a cartoon movie. (*A* – amplitude, *B* – phase). Data shown as red crosses are represent mean±SEM across participants. *A*) Amplitude results. Noise floor is given by the mean amplitude of the fitted pupil response to the *Reference* modulations, which contained no additional power at the stimulation frequency. *B*) Phase data, with negative angles corresponding to a phase delay (pupil response lags behind the stimulus), and positive angles corresponding to a phase advance (pupil response precedes the stimulus). The amplitude and phase data shown here correspond directly to the red fit lines in Fig. 2.

### Amplitude and phase of pupil responses

Expressed as amplitude and phase, we find that LMS and light flux pupil responses are well above the noise floor defined by the aggregate fit of the pupil response to the stimulus containing no stimulus at the modulation frequency (0.25 Hz). For all observers, the phase of the LMS and light flux response is delayed relative to the stimulus onset (Fig. 2A, C; 3B), which is consistent with the “dead time” of the human pupil response (∼200-300 msec). For the melanopsin-evoked pupil response, we find variability in both amplitude and phase (Fig. 2B; 3A, B), which warrants further investigation.

### Summation of pupil responses

We examined to what extent the sum of the responses to the *LMS* and *Mel* stimuli resembles the response to the *Light Flux* stimulus, which is, when expressed in contrast, the sum of the *LMS* and *Mel* stimuli in the stimulus domain: The *Light Flux* stimulus modulates the L, M, and S cones in the same way that the LMS modulation does; and it modulates melanopsin in the same way that the *Mel* modulation does. We applied a simple linear model by adding the amplitude and phase, expressed as complex sum, of the *LMS* and *Mel* responses, and reading out the amplitude again (Fig. 4). Conceptually, this is a vector sum model, where the pupil response to each modulation corresponds to a vector (defined by a given phase and amplitude), and we simply add vectors (Fig. 4, inset), yielding a new vector. We find that the amplitude of the complex sum of the *LMS* and *melanopsin* pupil response can account for the *Light Flux* pupil response while the *LMS* response only does so in a limited way (Fig. 4). We note that a better fit of the amplitude of the complex sum already requires that the phase of the two summands is aligned in such a way as to yield a better-fitting amplitude, we did not consider the goodness-of-fit of phase.

**Figure 4.**
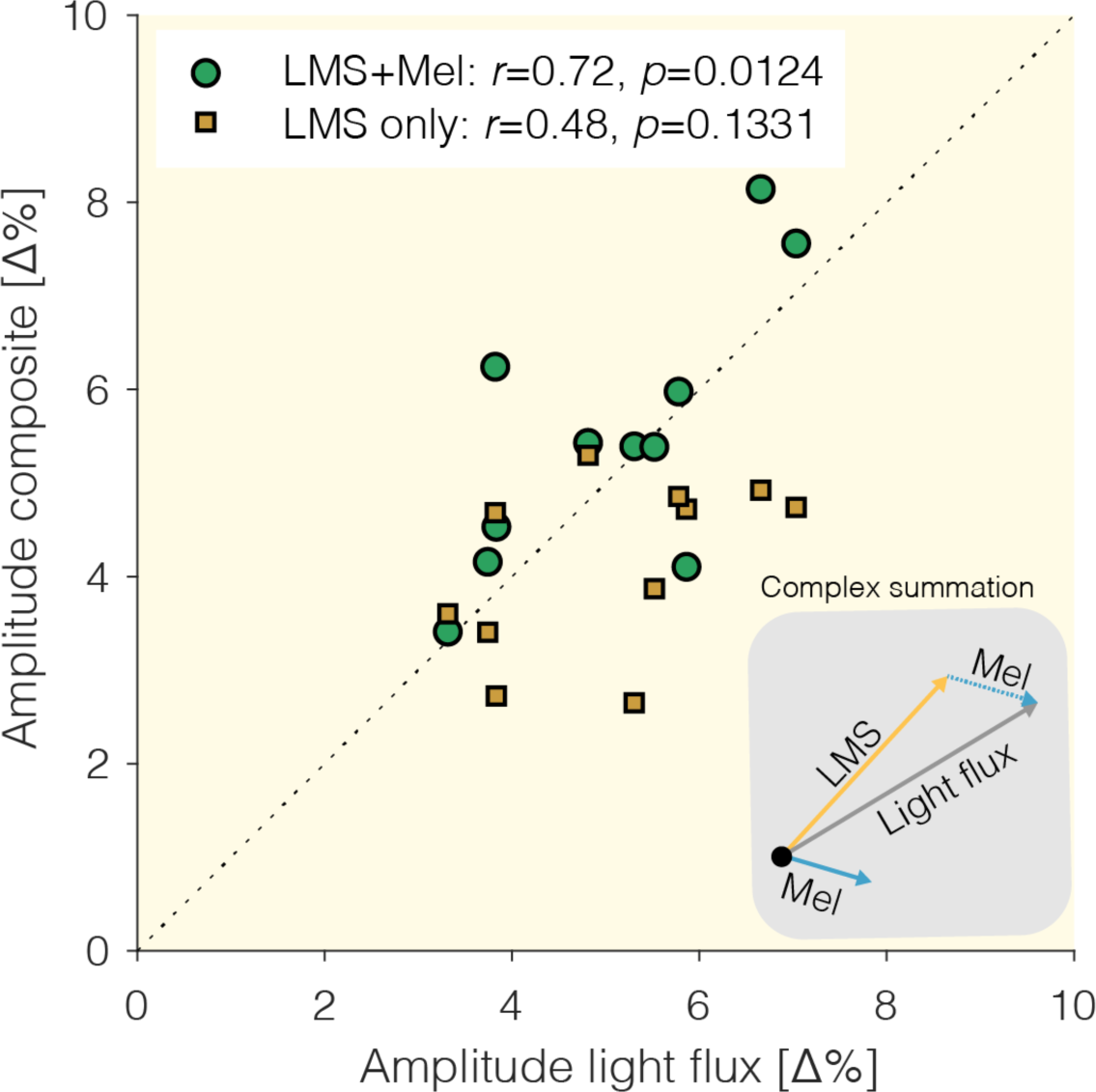
Linear summation model. We compare the amplitude of the pupil response to the *Light flux* modulation to the amplitude of two candidate models: The response to the *LMS* modulation, and the complex sum of the response to the *LMS* modulation and the response to the *Mel* modulation. *Inset*: Visualization of the vector summation model.

## Discussion

### Summary of results

Pupillary responses to melanopsin-directed stimuli using the method of silent substitution have been documented in the literature [7–16, 24], including by the authors [12, 24]. These studies typically have employed very carefully controlled artificial stimuli. However, past work has not addressed the question of how these photoreceptors each define pupil size during natural viewing. In this study, we find that by embedding photoreceptor-selective modulations in cartoon movie stimuli viewed under real-world conditions, we can elicit pupil responses specific to the receptor type. Consistent with what we would predict from studies employing spatially homogenous stimuli, we find a subtle but significant response to the melanopsin-mediated stimulus at the stimulation frequency (0.25 Hz). The response to the *Light flux* modulation is in amplitude larger than the response to the *LMS* modulation, providing additional evidence that there are melanopsin contributions to early visual processing that can be detected in our paradigm. Further, we find that we can model the pupil response to the *Light flux* modulation as the complex sum of the responses to the *Melanopsin* and the *LMS* modulation, which is corroborated by an earlier finding [12] (see their Fig. S8).

### Defining pupil size during free viewing of movie stimuli under real-world conditions

By embedding the modulations in an “entertainment” movie, we have placed the task in an ecologically-relevant context for using a video display unit (VDU). Allowing the participants to freely view the movie and not adhere to specific fixation procedures gives the paradigm an additional real-world aspect. While much emphasis has been placed on examining the photoreceptor inputs into pupillary control under well-controlled but relatively reduced stimulus conditions, our approach provides an understanding of these inputs that is complementary to more parametric and analytical approaches. Using this pragmatic approach to see if we could modulate the activity of photoreceptors in a task that human participants might engage in, in a normal day-to-day sense, we indeed confirm that all photoreceptors known to be active at photopic light levels (as established in previous studies, e.g. [12, 13]), can be manipulated using modulations embedded in movies which can be watched in real-world conditions.

### Uncontrolled factors

We wish to highlight that this study has purposely not accounted for a set of factors known to influence pupil size, on the grounds that the goal of this study was to examine pupil responses in an approximation of real-world conditions. Because of the near triad, vergence eye movements, changes of accommodation, and pupillary response go hand in hand when shifting view from near to far distances [33]. Under the unconstrained viewing conditions employed in this study, observers were able to move both head and eyes freely, thus introducing pupil changes in pupil diameter unrelated to our experimental parameter under control, photoreceptor contrast. As the stimuli were presented using a projection system in an otherwise dark room, eye movements of any sort could displace high contrast edges between the illuminated part of the canvas and the rest of the (dark) wall, which could lead to a transient signal. Because the modulation of contrast was uncorrelated with changes in movie content (and indeed, agnostic to it) which might have elicited gaze shifts, we feel confident that any effects of accommodation and vergence on pupil size average out, as evidence by the flat pupil response in the reference condition. Changes in autonomic arousal, which leads to differences in pupil size [34], might have occurred during the study, which was addressed by counterbalancing the order stimulus conditions.

### Data variability

We note that there appears to be large variability in the data. These could reflect individual differences, though this is not explicitly addressed in our data. The variability in the melanopsin response, both in amplitude and phase, warrants further investigation.

### Challenges to silent substitution

The silent substitution nominally allows selective stimulation of a given photoreceptor class. Silent substitution requires, however, a good estimate of the human participants’ spectral sensitivities. In practice, these are assumed based on standard cone fundamentals such as the physiologically-relevant CIE cone fundamentals [29]; some investigators also allow for an individual-observer calibration routine (e.g. [9–11, 13, 23]). In this work, for practical and pragmatic reasons, we forwent calibration at the level of the individual observer, and simply accounted for age-dependent differences in pre-receptoral filtering, which largely affects the amount of short-wavelength filtering, within a given age bracket spanning 5 years. In our modulations, we did not account for the potential intrusion of rods at daylight levels [35, 36]. We note that if rods were considered to contribute to these types of modulations, they would do so in a similar way for any practical application involving VDUs.

### Opportunities for translation

Given the broad impact of light on aspects of human physiology and behavior (such as melatonin suppression, circadian phase-shifting and modulation of alertness), the system and paradigm described here to modulate visual responses in a photoreceptor-selective fashion represents an opportunity to modulate these while people are engaging in everyday tasks such as movie-watching [17]. One powerful further application in the future could be as a method for low-impact “passive” assessment of photoreceptor health in clinical contexts, specifically in pediatric and geriatric populations.

### Towards a revision of colorimetry?

In this study, we have employed a 5-primary projector-based system for silent substitution with spatial control at the pixel level, and controlling the activation of the photoreceptors active at daylight: the cones and the melanopsin-containing ipRGCs. Commercially available VDUs have three primaries, which reflects the fact that human colour vision in photopic light levels is trichromatic. However, this study and a recent one [17] employing a similar system find that non-visual responses can be manipulated with a modified VDU with more than three primaries. Additionally, there is more and more evidence that melanopsin contributes to “classical” visual functions such as the perception of brightness [22, 37], colour [9, 38], space (humans: [9]; mice: [39]), and other visual attributes (humans: [24], mice: [40–42]). It remains an interesting question to what extent the classical colour matching functions, which are the basis of our current cone fundamentals, reflect the activity of melanopsin (though the amount of rod contributions have been discussed extensively [43]). Given the numerous roles of melanopsin in visual perception and in regulating physiology and behavior, and the emergence of tetrachromatic or higher-order primary displays, we expect that colorimetry may need some revisions to account for these effects, visual or non-visual.

## Acknowledgements

We wish to thank the EEG Lab at the University of Manchester for allowing us to use their space.

## References

1. Provencio, I., et al., A novel human opsin in the inner retina. J Neurosci, 2000. 20(2): p. 600–5.

2. Berson, D.M., F.A. Dunn, and M. Takao, Phototransduction by retinal ganglion cells that set the circadian clock. Science, 2002. 295(5557): p. 1070–3.

3. Hattar, S., et al., Melanopsin-containing retinal ganglion cells: architecture, projections, and intrinsic photosensitivity. Science, 2002. 295(5557): p. 1065–70.

4. Lucas, R.J., et al., Diminished pupillary light reflex at high irradiances in melanopsin-knockout mice. Science, 2003. 299(5604): p. 245–7.

5. Dacey, D.M., et al., Melanopsin-expressing ganglion cells in primate retina signal colour and irradiance and project to the LGN. Nature, 2005. 433(7027): p. 749–54.

6. Lucas, R.J., et al., Measuring and using light in the melanopsin age. Trends Neurosci, 2014. 37(1): p. 1–9.

7. Woelders, T., et al., Melanopsin- and L-cone-induced pupil constriction is inhibited by S- and M-cones in humans. Proc Natl Acad Sci U S A, 2018. 115(4): p. 792–797.

8. Murray, I.J., et al., Paradoxical pupil responses to isolated M-cone increments. J Opt Soc Am A Opt Image Sci Vis, 2018. 35(4): p. B66–B71.

9. Zele, A.J., et al., Melanopsin photoreception contributes to human visual detection, temporal and colour processing. Sci Rep, 2018. 8(1): p. 3842.

10. Barrionuevo, P.A. and D. Cao, Luminance and chromatic signals interact differently with melanopsin activation to control the pupil light response. J Vis, 2016. 16(11): p.29.

11. Cao, D., N. Nicandro, and P.A. Barrionuevo, A five-primary photostimulator suitable for studying intrinsically photosensitive retinal ganglion cell functions in humans. J Vis, 2015. 15(1): p. 15 1 27.

12. Spitschan, M., et al., Opponent melanopsin and S-cone signals in the human pupillary light response. Proc Natl Acad Sci U S A, 2014. 111(43): p. 15568–72.

13. Barrionuevo, P.A., et al., Assessing rod, cone, and melanopsin contributions to human pupil flicker responses. Invest Ophthalmol Vis Sci, 2014. 55(2): p. 719–27.

14. Tsujimura, S. and Y. Tokuda, Delayed response of human melanopsin retinal ganglion cells on the pupillary light reflex. Ophthalmic Physiol Opt, 2011. 31(5): p. 469–79.

15. Vienot, F., S. Bailacq, and J.L. Rohellec, The effect of controlled photopigment excitations on pupil aperture. Ophthalmic Physiol Opt, 2010. 30(5): p. 484–91.

16. Tsujimura, S., et al., Contribution of human melanopsin retinal ganglion cells to steady-state pupil responses. Proc Biol Sci, 2010. 277(1693): p. 2485–92.

17. Allen, A.E., et al., Exploiting metamerism to regulate the impact of a visual display on alertness and melatonin suppression independent of visual appearance. Sleep, 2018. 41(8).

18. Spitschan, M., G.K. Aguirre, and D.H. Brainard, Selective stimulation of penumbral cones reveals perception in the shadow of retinal blood vessels. PLoS One, 2015. 10(4): p. e0124328.

19. Estévez, O. and H. Spekreijse, The “silent substitution” method in visual research. Vision Res, 1982. 22(6): p. 681–91.

20. Estévez, O. and H. Spekreijse, A spectral compensation method for determining the flicker characteristics of the human colour mechanisms. Vision Res, 1974. 14(9): p. 823–30.

21. Vartanian, G., K.Y. Wong, and P.C. Ku, LED Lights With Hidden Intensity-Modulated Blue Channels Aiming for Enhanced Subconscious Visual Responses. IEEE Photonics J, 2017. 9(3).

22. Brown, T.M., et al., Melanopsin-based brightness discrimination in mice and humans. Curr Biol, 2012. 22(12): p. 1134–41.

23. Pokorny, J., H. Smithson, and J. Quinlan, Photostimulator allowing independent control of rods and the three cone types. Vis Neurosci, 2004. 21(3): p. 263–7.

24. Spitschan, M., et al., The human visual cortex response to melanopsin-directed stimulation is accompanied by a distinct perceptual experience. Proc Natl Acad Sci U S A, 2017. 114(46): p. 12291–12296.

25. Spitschan, M., et al., Human Visual Cortex Responses to Rapid Cone and Melanopsin-Directed Flicker. J Neurosci, 2016. 36(5): p. 1471–82.

26. MacKinnon, N., et al., Spectrally programmable light engine for in vitro or in vivo molecular imaging and spectroscopy. Appl Opt, 2005. 44(11): p. 2033–40.

27. Bayer, F.S., et al., A tetrachromatic display for the spatiotemporal control of rod and cone stimulation. J Vis, 2015. 15(11): p. 15.

28. Tom And Jerry - Complete Volumes 1–6. 2006, Warner Home Video.

29. CIE, Fundamental chromaticity diagram with physiological axes – Part 1 (Technical Report 170-1). 2006, Vienna: Central Bureau of the Commission Internationale de l’ Éclairage.

30. Xu, J., J. Pokorny, and V.C. Smith, Optical density of the human lens. J Opt Soc Am A Opt Image Sci Vis, 1997. 14(5): p. 953–60.

31. Pokorny, J., V.C. Smith, and M. Lutze, Aging of the human lens. Appl Opt, 1987. 26(8): p. 1437–40.

32. Ishihara, S., *Tests for Colour Blindness*. 1977, Tokyo: Kanehara Shuppen Company.

33. McDougal, D.H. and P.D. Gamlin, Autonomic control of the eye. Compr Physiol, 2015. 5(1): p. 439–73.

34. Loewenfeld, I.E. and O. Lowenstein, The pupil: anatomy, physiology, and clinical applications. 1st ed. 1993, Ames / Detroit: Iowa State University Press / Wayne State University Press.

35. Tikidji-Hamburyan, A., et al., Rods progressively escape saturation to drive visual responses in daylight conditions. Nat Commun, 2017. 8(1): p. 1813.

36. Shapiro, A.G., Cone-specific mediation of rod sensitivity in trichromatic observers. Invest Ophthalmol Vis Sci, 2002. 43(3): p. 898–905.

37. Zele, A.J., et al., Cone and melanopsin contributions to human brightness estimation. J Opt Soc Am A Opt Image Sci Vis, 2018. 35(4): p. B19–B25.

38. Cao, D., A. Chang, and S. Gai, Evidence for an impact of melanopsin activation on unique white perception. J Opt Soc Am A Opt Image Sci Vis, 2018. 35(4): p. B287–B291.

39. Allen, A.E., et al., Melanopsin Contributions to the Representation of Images in the Early Visual System. Curr Biol, 2017. 27(11): p. 1623–1632 e4.

40. Allen, A.E., et al., Melanopsin-driven light adaptation in mouse vision. Curr Biol, 2014. 24(21): p. 2481–90.

41. Storchi, R., et al., Melanopsin-driven increases in maintained activity enhance thalamic visual response reliability across a simulated dawn. Proc Natl Acad Sci U S A, 2015. 112(42): p. E5734–43.

42. Storchi, R., et al., Modulation of Fast Narrowband Oscillations in the Mouse Retina and dLGN According to Background Light Intensity. Neuron, 2017. 93(2): p. 299–307.

43. Wyszecki, G. and W.S. Stiles, Color Science: concepts and methods, quantitative data and formulæ. 2nd ed. 1982, New York: Wiley.

